# Soda-baited RNAi Yeast Insecticides as Effective Attractive Targeted Sugar Baits (ATSBs) for Mosquito Control

**DOI:** 10.64898/2026.05.01.722182

**Authors:** Akilah T. M. Stewart, Majidah Hamid-Adiamoh, Longhua Sun, Teresia M. Njoroge, Nikhella Winter-Reece, Rachel Shui Feng, Satish Singh, Lester D. James, Britton Sofhauser, Carolina Dille, Azad Mohammed, David W. Severson, Molly Duman-Scheel

**Affiliations:** Department of Medical and Molecular Genetics, Indiana University School of Medicine, Raclin-Carmichael Hall, South Bend, Indiana, United States of America; Eck Institute for Global Health, University of Notre Dame, Notre Dame, Indiana, United States of America; Department of Life Sciences, Faculty of Science and Technology, The University of the West Indies, St. Augustine Campus, Trinidad and Tobago; College of Arts and Letters, The University of Notre Dame, Notre Dame, Indiana, United States of America; Department of Biological Sciences, The University of Notre Dame, Notre Dame, Indiana, United States of America

**Keywords:** *Aedes*, *Anopheles*, attractive targeted sugar bait (ATSB), *Culex*, mosquito, mosquito control, RNA interference (RNAi), semi-field trial, sugar baits, sugar feeding behavior, vector control, yeast

## Abstract

**Background:** Attractive Targeted Sugar Baits (ATSBs) are a promising, environmentally friendly approach for mosquito control, but the direct field application, scalability and long-term effectiveness of ATSBs across diverse mosquito species remain significant challenges.

**Methodology/Principal Findings:** We assessed the efficacy of a genetically engineered RNA interference (RNAi) yeast strain (Sh.463_56.10R) formulated in three sugar baits, soda (Coca-Cola^TM^), 10% sucrose, and a commercial mosquito bait (BaitStab^TM^), on *Aedes, Anopheles* and *Culex* mosquitoes. All RNAi yeast bait formulations induced significantly higher mean mortality (87-100%) compared to the control groups (0-9%; *P*<0.0001), but mosquitoes exhibited a higher feeding preference for RNAi yeast-soda baits, which induced mortality rates of 94-100% (*P* < 0.0001) recorded across all mosquito species. Additionally, to assess the competitiveness of the RNAi yeast-soda bait to other tropical sugar sources, semi-field choice assays were conducted in Trinidad, West Indies using competing flowering plants and fruits typically found in residential environments. The RNAi yeast-soda ATSB continued to perform well in the presence of competing floral and fruit sugar sources during both *Aedes albopictus* and *Culex quiquefasciatus* trials, though the presence of several fruits and flowers did reduce *A. aegypti* mortality, suggesting that further field testing will be necessary. The residual activity of the Sh.463_56.10R + soda formulation was retained for at least 14 months, with sustained 100% mortality in *C. quinquefasciatus* and 93-100% mortality in *Aedes aegypti*, *Anopheles gambiae* and *Anopheles stephensi*. The RNAi yeast-soda ATSB also performed well in semi-field studies performed with a prototype soda bottle feeder.

**Conclusions/Significance:** This study demonstrates the potential of soda-baited RNAi yeast as a potent, long-lasting, and scalable platform for ATSB-based mosquito control as a component of integrated vector management programs.

**Author Summary:** Mosquito-borne diseases continue to affect millions of people worldwide, and current mosquito control methods face challenges such as low public uptake, insecticide resistance and environmental concerns. Here we evaluated a new and environmentally friendly approach to mosquito control using ATSBs. We tested genetically engineered species-specific yeast producing RNAi molecules capable of killing mosquitoes that feed on it. We mixed the yeast with three different sugar baits, including soda (Coca-Cola^TM^), 10% sucrose, and the commercial mosquito bait BaitStab^TM^ formulation, and evaluated how well they worked against different mosquitoes. The results showed that the RNAi yeast mixed with soda was the most effective, killing up to 100% of mosquitoes in laboratory and outdoor tests. The bait remained effective in the presence of many competing natural tropical fruit and floral sugar sources. Remarkably, the bait, which can be delivered in a soda bottle feeder, stayed active for at least 14 months under simulated field conditions. These findings suggest that soda-based RNAi yeast baits could provide a practical, long-lasting and scalable tool for mosquito control and may help strengthen future strategies to reduce mosquito-borne diseases.

## Introduction

Mosquito-borne diseases pose significant threats to global health, accounting for nearly one million deaths annually worldwide (1). The primary strategy for preventing these diseases is mosquito control using synthetic insecticides. However, the long-term effectiveness of this approach is challenged by low adherence to control measures, increasing instances of insecticide resistance, and growing concerns about the harmful effects of insecticides on non-target organisms (2, 3). Addressing these challenges is crucial for ensuring the effectiveness and sustainability of the current strategies. The success of the World Health Organization (WHO) global plan for insecticide resistance management (4) relies on the effective implementation of new insecticides (3). Hence, extensive research efforts are underway to develop novel active ingredients with alternative modes of action, as well as innovative approaches such as biological control, genetic modification, and behaviorally based interventions to reduce reliance on chemical insecticides and enhance the sustainability of vector control programs (5).

RNA interference (RNAi) is an emerging and promising approach with significant potential for mosquito control. Our recent studies (6–11) have demonstrated that RNAi pesticides, which can be inexpensively produced in yeast, can silence genes required for mosquito survival. These novel pesticides are designed specifically to target conserved sequences in mosquitoes that are not found in non-target organisms. The pesticides are delivered as oral insecticides using heat-inactivated lyophilized yeast as a carrier, and the lethal yeast is capable of effectively killing mosquito vectors, including *Aedes*, *Culex* and *Anopheles spp.,* without any impact on non-target organisms (6–11).

ATSBs, a new paradigm for vector control, capitalize on the natural sugar feeding behavior of female and male mosquitoes that are drawn to feed on a sugar bait mixed with an insecticide. ATSB-mediated delivery of several non-specific insecticides has been successfully applied for direct control of *A. aegypti*, *A. albopictus* and other *Aedes* species (12–19) both in laboratory and field conditions. Similarly, *Culex pipiens, C. quinquefasciatus, C. tarsalis* and a variety of other *Culex* species, including nuisance biters, have been successfully targeted with ATSBs (13, 20–23). In addition, ATSBs are also showing great promise for targeting *Anopheles* species (24–26). In field trials, ATSBs reduced the density of *Anopheles* mosquitoes (24–26). However, the evolution of resistance mechanisms remains a significant challenge for the use of ATSBs, underscoring the need to identify second-generation ATSB insecticides. Moreover, despite the addition of protective barriers to bait stations and efforts to limit ATSB applications to non-flowering vegetation, it is difficult to completely eliminate risks to pollinators and other non-target organisms (27).

Given that the success of ATSB deployment relies on effectively diverting mosquitoes from natural sugar sources, it is critical that baits which are highly attractive to mosquitoes are used. We have developed a novel mosquito-specific yeast insecticide, Sh.463 (8), that expresses interfering RNA which targets a conserved site in the *Shaker (Sh)* gene of multiple species of mosquitoes, but not humans or other non-target organisms, including honeybees. This eco-friendly universal mosquito insecticide has performed well in lab and semi-field trials (6, 8, 28), promoting increased sugar feeding rates and killing up to 100% of mosquitoes when deployed using the commercial BaitStab^TM^ matrix. However, when ATSBs prepared with BaitStab^TM^ and the broad-based chemical insecticide dinotefuran were evaluated in recent Phase III clinical trials (29), despite the initially promising entomological impacts of these ATSB baits (26), significant epidemiological impacts were ultimately not observed (29). It is also presently unclear if the baits deployed in the trial, which assessed malaria transmission, will be the most effective lure for *Aedes* and *Culex* arboviral vector mosquitoes in the field.

Building on our initial data from the Sh.463 (8) RNAi yeast, the addition of which may increase the attractiveness of BaitStab^TM^ (26) in both laboratory (6) and simulated semi-field conditions (28), this study aimed to further enhance the effectiveness of the yeast by identifying a more attractive sugar bait formulation for use in Sh.463-based ATSB stations targeting multiple mosquito species. We hypothesized that incorporating soda-based sugar formulations with the Sh.463 RNAi yeast insecticide will significantly increase mosquito feeding preference and mortality across multiple mosquito species, thereby enhancing the efficacy of ATSBs for mosquito control. This study assessed the feeding preference using mortality as an indicator across different combinations of Sh.463 RNAi yeast with three sugar baits in *Aedes, Anopheles* and *Culex* mosquitoes. We further assessed the ATSB efficacy in semi-field conditions and against natural plant sugar sources available to mosquitoes in tropical environments.

## Methods

### Preparation of RNAi yeast

A previously developed broad-based, high-expression Sh.463 RNAi yeast strain (6) that expresses short hairpin RNA (shRNA) targeting conserved sequences in mosquito *Sh* genes across multiple mosquito species (8) was used in this investigation. The genotype of the Sh.463 yeast strain (hereafter referred to as Sh.463_56.10R) is: *MATa*, *PiggyBac* (*leu2d/P_TDH3_-shRNA_463-T_CYC1_*, *P_TDH3_-shRNA_463-T_CYC1_*, *P_TDH3_-shRNA_463-T_CYC1_*), *CEN/ARS* (*URA3/SPBase-Sc-CO*) (6). The control strain (hereafter referred to as Control_347.1R), which expresses a hairpin with no known target in any mosquito species, has the following genotype: *MATa, PiggyBac* (*LEU2/P_TDH3_-shRNA_Ctrl-T_CYC1_*), *2um* (*URA3/SPBase_Sc-CO*), *PiggyBac* (*HIS3/P_TDH3_-shRNA_Ctrl-T_CYC1_*), *CEN/ARS* (*URA3/SPBase_Sc-CO*) *PiggyBac* (*trp1d/P_TDH3_-shRNA_Ctrl-T_CYC1_*), *CEN/ARS* (*URA3/SPBase_Sc-CO*) (6). The yeast was prepared, heat killed and lyophilized as described (30).

### Mosquito rearing for laboratory experiments

The mosquito strains used for all laboratory experiments included *A. albopictus* Gainesville (MRA-804, Biological and Emerging Infections (BEI) Research Resources Program, NIAID, NIH*), A. aegypti* Liverpool-IB12 (LVP-IB12), *A. gambiae* G3 (BEI Resources: MRA-112, contributed by Mark Q. Benedict), *A. stephensi* strain STE2 (MRA-128, BEI Resources), *C. quinquefasciatus* JHB (NR-43025, BEI Resources), and *C. pipiens* (developed from Niles, MI, USA ovitrap collection) (9). Mosquitoes were reared in the insectary facility at the Indiana University School of Medicine, United States, under standard insectary conditions (31): 27 ± 2 °C; 70 ± 10% relative humidity and 12-h light/dark cycle.

### Preparation of sugar formulations

Three sugar formulations were tested: 10% sucrose solution, BaitStab^TM^ matrix without dinotefuran (Westham Co., Israel) and soda (Coca-Cola^TM^), each combined with either the Sh.463_56.10R active ingredient (Treatment) or Control_347.1R RNAi yeast (Control). The 10% sucrose solution (ASB) represents the standard sugar for maintaining mosquito colonies and was prepared by mixing 10 g of sucrose with 100 ml of distilled water. The BaitStab^TM^ matrix [hereafter referred to as Westham bait (WH), acquired from Westham Co., Israel] is a proprietary sugar attractant (32) used in several multi-country ATSB field trials, with its effectiveness across these trials recently summarized (33). In the current study, the bait was used without any dinotefuran. Aliquots of this bait were used directly without any modification to the original formulation. Coca-Cola^TM^, which contains approximately 11% sucrose, was selected as the soda-based attractive sugar bait (ASB) due to its high sugar content and proven efficacy in inducing significant mosquito feeding (34), including with *Aedes japonicus* (35). Moreover, it is universally available and easily accessible as both an attractant and a sugar source for an ATSB. Apart from being degassed, the soda ASB was used in its original form without modification. Degassing was achieved by vigorously shaking a Coca-Cola^TM^ bottle through ten cycles, with the gas released by gently opening the lid at the end of each cycle. This degassing was critical for utility of the bait in the prototype feeder described (see below), as the feeder would leak when soda that had not been degassed was used.

### Mosquito yeast feeding experiments

Different experimental approaches were employed to assess mosquito feeding preferences using mortality rates and residual efficacy of the RNAi yeast in the three sugar baits. In all experiments except the fruit/flower competition assays discussed below, 25 non-blood-fed female mosquitoes (4-5 days old) were used. Prior to RNAi yeast feeding, mosquitoes were either sugar-starved for 24 hours (*A. gambiae)* or 48 h (*A. albopictus, A. aegypti, C. quinquefasciatus* and *C. pipiens*). Water was provided using cotton balls during starvation only for *A. gambiae* and *A. stephensi* (for residual experiments only). Mosquitoes were primarily fed with the RNAi yeast mixture (Treatment_Sh.463_56.10R/Control_347.1R + sugar bait) overnight, after which they were maintained on a 10% sucrose solution for six days when the trial was concluded. Mortality was monitored for six days following feeding. A total of nine biological replicates (three replicates per trial) were conducted for each sugar bait experiment.

All experiments performed with the lethal yeast Sh.463_56.10R (Treatment) strains were designated as ATSBs or Treatment, whereas those using Control_347.1R yeast were referred to as Control. In addition, experiments conducted with the individual sugar formulations alone, categorized as Attractive Sugar Baits or ASBs, were also performed. The combinations of RNAi yeast and sugar formulations used in all experiments are henceforth identified as follows:

i. Treatment-Sugar: Sh.463_56.10R + 10% sugar
ii. Control-Sugar: Control_347.1R + 10% sugar
iii. ASB-Sugar: 10% sugar
iv. Treatment-WH: Sh.463_56.10R + WH
v. Control-WH: Control_347.1R + WH
vi. ASB-WH: WH
vii. Treatment-Soda: Sh.463_56.10R + soda
viii. Control-Soda: Control_347.1R + soda
ix. ASB-Soda: Coca-Cola^TM^

### Evaluation of mosquito mortality in the three RNAi yeast-sugar formulations

ATSB trials were conducted with individual sugar formulations inside circular (3.7L) mosquito cages mainly following the previously described protocol (7, 8, 10, 11). Briefly, a paste of sugar bait-yeast mixture containing 40 mg of lyophilized yeast and 120 μL of sugar bait (final concentration: 333.3 μg/μL), was prepared and fed to starved mosquitoes inside cages. Following initial optimization, the Treatment-Sugar and ASB-Sugar experiments were presented to mosquitoes in petri dishes, whereas the Treatment-WH and ASB-WH experiments were presented in a sealed miniature sachet feeder made with 5 and 25 micron food-grade nylon membranes (Amazon, Seattle Washington) and prepared as described (9). The baits were administered using MUDUODUO automatic bird drinker cups (Amazon, Seattle Washington) custom-designed as ATSB feeders as described (36). Briefly, a yeast paste containing 200 mg of either control or treatment yeast in 300-360 µL of soda (556-667 µg/µL) was evenly spread inside a sachet made from the same food-grade nylon membrane used above. The sachet was then placed in the well of a MUDUODUO automatic bird feeder cup. A 12 oz Coca-Cola^TM^ bottle was filled with 120 mL of degassed soda and inverted onto the feeder to provide a continuous supply of soda to the yeast inside the sachet. The membrane was secured in place using a wick made from a small piece of clean dehumidifier filter (Honeywell Home, Charlotte, NC). Mortality was recorded for six days post-yeast feeding.

### Comparative assessment of mosquito feeding preference for soda versus alternative sugar baits

Dual-choice ATSB experiments were conducted to evaluate mosquito feeding preferences between different sugar bait formulations. The experiments were performed in 30 cm × 30 cm x 30 cm Bug Dorm cages (MegaView Science, Taiwan) using both Treatment ATSBs and the corresponding Control for each sugar formulation simultaneously. In each experimental cage, two baits were presented: one containing the Treatment yeast Sh.463_56.10R and the other containing Control yeast 347.1R in either of two sugar formulations. ASBs for each sugar type were also included in separate negative control cages.

Specifically, for the dual-choice experiments between soda and 10% sugar ASBs, Treatment-Soda (Sh.463_56.10R + soda) and Control-Sugar (Control_347.1R + 10% sugar) were placed in one cage, while Control-Soda (Control_347.1R + soda) and Treatment-Sugar (Sh.463_56.10R + 10% sugar) were placed in a separate cage. Similarly, in the soda versus WH comparison, the bait pairs included Treatment-Soda-and Control-WH in one cage, and Control-Soda and Treatment-WH in another.

All baits were positioned in opposite corners of the experimental cages to ensure equal exposure. To reduce positional access bias, the locations of the baits were rotated in each replicate. The RNAi yeast-sugar mixtures were prepared at the same final concentration of 333.3 μg/μL as described above but at double the volume and applied according to the optimal delivery method for each formulation as described above. Mosquitoes were fed overnight in the soda versus sugar trials, and for 4 hours in the soda versus WH trials, based on optimized feeding durations. Each dual-choice pair was tested in three technical replicates, and mortality was also monitored for six days.

### Semi-field assessment using RNAi yeast in soda feeders

To assess the most optimal bait from previous trials in ambient environmental conditions, experiments were also conducted in semi-field conditions following the Yeast-ATSB-Soda and ASB-Soda experiments described above and as previously reported (36). The experiment was performed using *C. pipiens* females in September and October 2025 at the Indiana University School of Medicine. Three biological trials, each consisting of three technical replicates, were performed. The same feeders were used for all the trials over a four-week period. During the semi-field trial period (n = 27 days), the average daily temperature was 20.6 ± 5.2 °C and ranged from lows of 6.1 °C to highs of 41.1 °C. The average percentage relative humidity was 45.4 ± 15.2% and ranged from 25.5 to 93.8%. The average daily minimum percentage relative humidity was 45.7 ± 13.2%, and the average daily maximum percentage relative humidity was 79.5 ± 10.0%.

### Semi-field evaluation of mosquito attraction to RNAi yeast-soda formulations versus natural plant sugar sources

Semi-field-based dual choice experiments were also conducted to determine whether mosquitoes demonstrate a feeding preference for soda-based RNAi yeast ATSB (Coca-Cola^TM^) over sugar sources derived from local fruits and flowers. These experiments were carried out between May and August 2023 at the rooftop laboratory of the University of the West Indies, St. Augustine Campus in Trinidad, West Indies. The experimental flowers and fruits tested included *Allamanda cathartica* (yellow trumpet), *Catharanthus roseus* (Madagascar periwinkle), *Hibiscus rosa-sinensis* (tropical hibiscus), *Ixora coccinea* (Jungle geranium), and *Lantana camara* (West Indian lantana), while the fruits tested were *Carica papaya* (papaya) and *Mangifera indica* (mango), all of which were locally sourced in Trinidad. These flowers and fruits were selected based on their local availability and abundance in nurseries, typical households and backyards in Trinidad. Flower cuttings were kept in water to maintain freshness; however, parafilm was used to prevent mosquito access to the water source and accidental drowning. The sugar sources from these natural plants were compared against soda combined with RNAi yeast. For these trials 80 mg of Sh.463_56.10R yeast was mixed with 200 μL of soda. The mixture was placed in 2cm x 4cm sachets made from unstretched Parafilm® “M” Laboratory Film (Amcor, Switzerland), and 12 pin sized holes were pierced for mosquito access. Sachets were vertically mounted in the cage, and no additional sugar solution was provided during the experiment.

Experiments were conducted using 15 female and 15 male adult mosquitoes in each 30 cm × 30 cm × 20 cm cage. 4-5 day old adult mosquitoes were starved for 24 h prior to the trials. The mosquito species used included *A. aegypti*, *A. albopictus*, and *C. quinquefasciatus*, all reared from at least the second generation of field-collected specimens gathered locally in ambient conditions at the university insectary. Each set of experiments included three parallel cage trials in which mosquitoes were exposed to sugar as follows:

1. A fruit or flower alone (with survival in these cages demonstrating that the mosquitoes could feed on it, which was required for survival during the six-day trial period).
2. The same fruit or flower as well as a sachet bait station consisting of a paste of Treatment-Soda (Sh.463_56.10R + soda), also referred to as a “Choice.”
3. Treatment-Soda paste alone, presented as a sachet and serving as a positive control.

The fruits/flowers and baits remained in the cage throughout the duration of the trial without additional sugar supplementation. Humidity was maintained by providing water via a saturated cotton ball in each cage. Each experiment included three biological replicates, with daily mortality recorded over six days. Temperatures ranged from 26.0 °C to 37.0 °C, and relative humidity varied between 34.0% and 90.0% during the experimental periods.

### Assessment of the residual activity of Sh.463_56.10R-Soda ATSB

The residual activity of the Sh.463_56.10R RNAi yeast as an insecticide in soda was evaluated over a fourteen-month period under insectary conditions. RNAi-ATSB mixtures were prepared at a final concentration of 333.3 μg/μL and delivered to mosquitoes in feeders, but with the soda bottle removed and the opening covered. The soda was continuously supplied by adding additional Coca-Cola^TM^ throughout the experimental period as needed. To prevent mold growth 100g of methyl-p-hydroxybenzoate (Carolina Biological Supply, Burlington, NC) was added per liter of 70% ethanol and added to degassed Coca-Cola^TM^ to a final concentration of 0.002%. Control baits (Control_347.1R + soda) and ASBs (soda with mold inhibitor only) were prepared similarly and included in all experiments.

In an effort to better simulate field conditions, all ATSBs were stored in the insectary for the duration of the experiments, which occurred over the course of 14 months. For each trial, 25 non-blood-fed female mosquitoes were fed overnight with the stored baits, and mortality was recorded daily for six days post-feeding. Fresh soda-mold inhibitor solution was added to replenish the feeder cup whenever drying was observed. The experiment was conducted on four mosquito species: *A. aegypti*, *C. quinquefasciatus* (25 mosquitoes per cage for each,) *A. gambiae* and *A. stephensi* (30 mosquitoes per cage for each). At least two biological replicates were performed per species per time point.

### Statistical analysis

Daily mortality data for each mosquito species were pooled and calculated as the percentage of mosquitoes that died relative to the total number exposed per experimental group. The mortality rates between RNAi yeast treatment and control yeast and ASB alone were not normally distributed (even following arcsine transformation), and so Kruskal-Wallis tests were performed followed by Dunn’s post hoc tests. Mosquito mortality rate was used as the metric to determine preference for each respective bait. Mann Whitney U tests were used to assess statistical differences between the percentages of mortality for the different sugar formulations in each experimental pair.

For semi-field choice experiments, Fisher’s Exact tests were used to assess mortality. One-way ANOVA followed by a post-hoc Tukey test HSD was conducted on the semi-field soda bottle feeder experimental data. For all analyses, a significance level of *P* < 0.05 was considered statistically significant. Statistical tests were performed using Statistical Package for the Social Sciences (SPSS) software Version 31 (IBM, Armonk, NY, USA).

### Institutional Review Board Statement

This study was approved by the Institutional Review Board of the University of the West Indies (UWI; Study CEC403/12/17), the Southwest Regional Health Authority, which is a division of the Trinidad and Tobago Ministry of Health, and by the head of the UWI St. Augustine Department of Life Sciences.

## Results

### Efficacy of RNAi yeast treatment across distinct ASB formulations

In laboratory trials, the evaluation of mosquito mortality using Sh.463_56.10R yeast (Treatment) as the active ingredient presented in three different baits: 10% sucrose (Sugar), WH bait (WH), and Coca-cola^TM^ (Soda) demonstrated the efficacy of the RNAi yeast in inducing high mortality in a broad range of mosquito species regardless of the sugar bait used (Fig. 1, Table S1). In all cases, mortality was significantly higher in the Treatment compared to either of the negative controls, which included Control_347.1R RNAi yeast (Control) or the sugar bait alone (ASB); significant differences in mortality rates were observed at *P*<0.001 in all species (Fig. 1, Table S1; except *P*<0.004 for ASB-Sugar on *C. quinquefasciatus*). For the Sh.463_56.10R + soda (Treatment-Soda) formulation, mortality ranged from 94.2 ± 2.54% (*C. pipiens*) to 100.0 ± 0.0% (*A. gambiae*) across all mosquito species. Mortality levels in the ASB and Control trials were negligible for all bait types across the species (Fig 1a).

**Fig 1.**
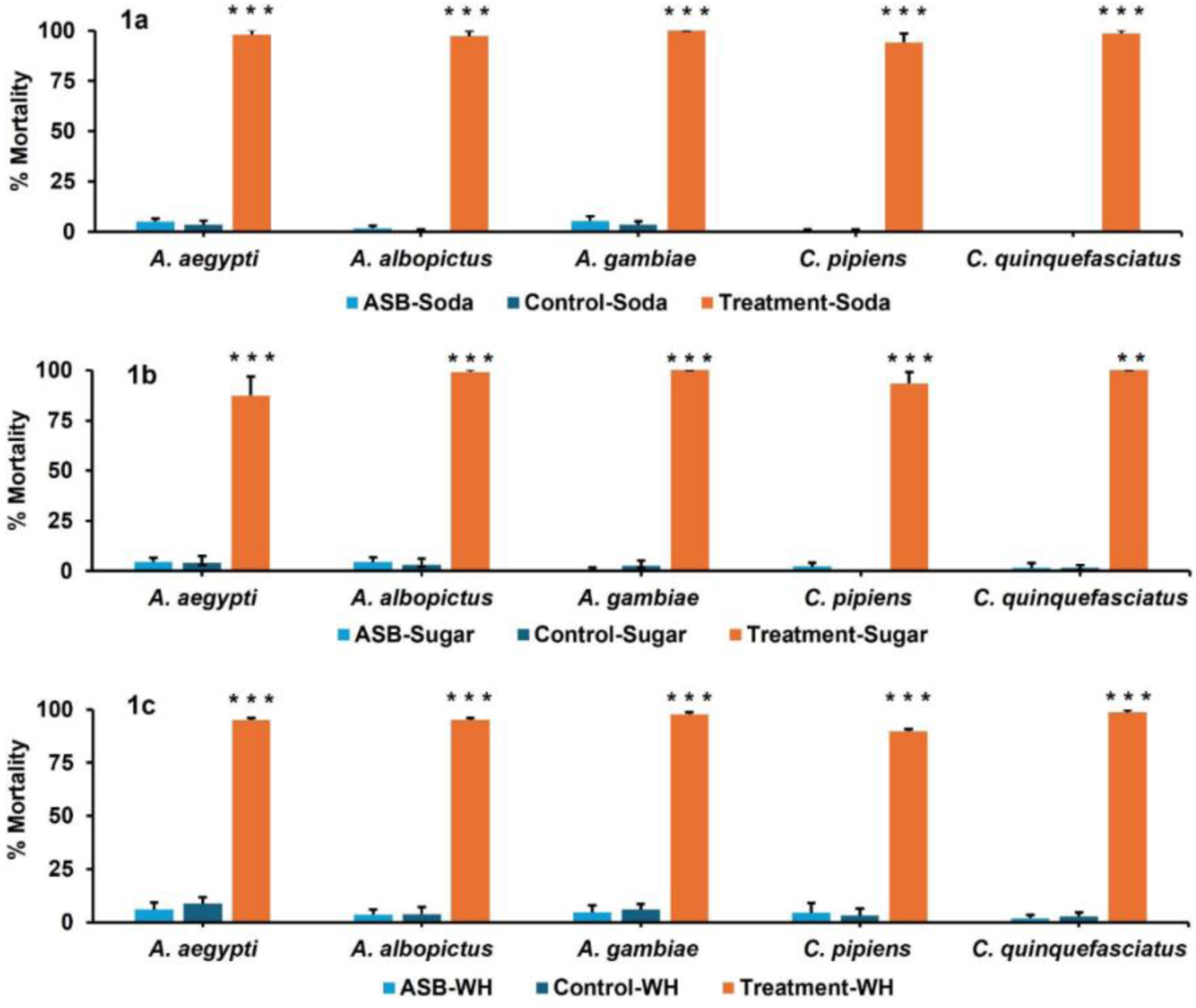
Mortality of the mosquito species exposed to three Sh.463_56.10R RNAi yeast treatment formulations. Mosquito mortality profiles in laboratory trials (Fig 1a: Soda, Fig 1b: Sugar, and Fig 1c: WH). Bars represent mean percentages of mortality across the five mosquito species exposed to the three treatments. *** (*P*≤0.001) or ** (*P*≤0.01) denote statistically significant differences in mortality for the Sh.463_56.10R treatment with respect to the ASB or Control for each species (Kruskal-Wallis Test with Dunn’s post hoc test), and error bars indicate standard error of the mean (SEM).

Similarly, for the 10% sugar bait (Sugar), Treatment-Sugar ATSBs induced high mortality across all mosquito species tested (Fig 1b). As with the soda formulation, *A. gambiae* and *C. quinquefasciatus* were completely killed (100%) in the Treatment-Sugar group. Mortality in *A. albopictus* (99.1 ± 0.59%) and *C. pipiens* (93.3 ± 3.40%) also exceeded 90%, whereas mortality in *A. aegypti* was 87.4 ± 6.20%. The mortality levels in the Treatment-Sugar cages were significantly more compared to the Control or ASB group mortalities (*P*<0.001).

The WH sugar bait formulation trials (WH) showed a similar pattern in which mortality was significantly higher (*P<*0.001) in Treatment-WH compared to the Control-WH or ASB-WH in all species tested (Fig 1c, S1 Table). Mortality in the Treatment-WH group ranged from 89.8 ± 3.31% *(C. pipiens)* to 98.7 ± 0.67% (*C. quinquefasciatus*).

### A mosquito feeding preference for soda ATSBs

Dual-choice experiments demonstrated that the RNAi yeast-soda formulation was the most attractive to all mosquito species compared to other sugar formulations. This preference was demonstrated through trials in which mosquitoes were given a choice between Treatment-Soda and Control-Sugar in one cage. Control-Soda and Treatment-Sugar selections were placed in a separate cage. Given the choices, all species exhibited a consistent feeding preference for Soda regardless of whether the Soda contained Treatment or Control yeast or its position within the cage, thus confirming the attractiveness of the RNAi yeast-soda ATSB formulation (*P*<0.01; Fig 2a; S2 Table).

**Fig 2.**
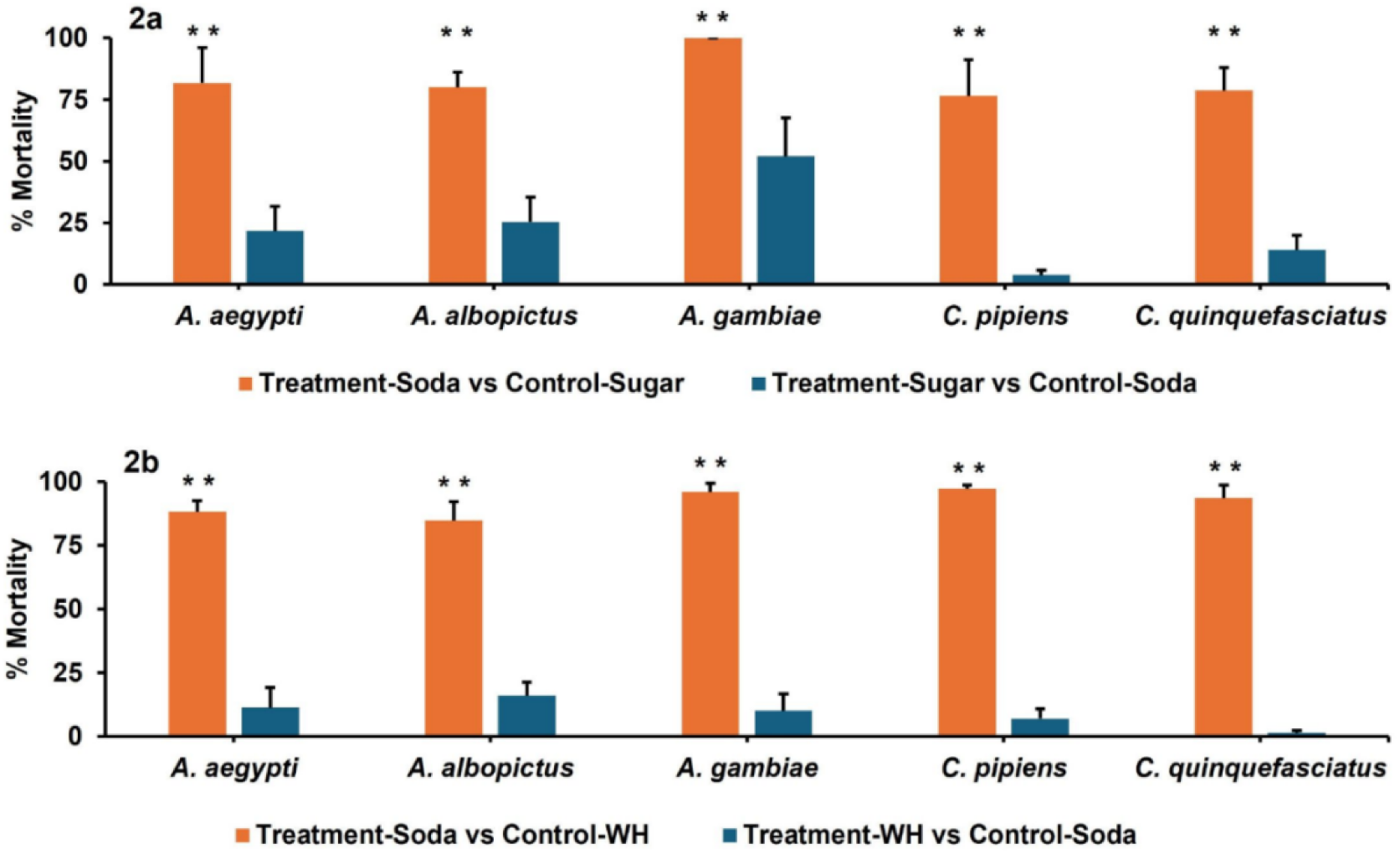
Dual choice experiments demonstrate a preference for yeast-soda baits. Shown are Treatment cage results from dual choice assays performed with: (2a): Treatment-Soda vs Control-Sugar (orange) or Treatment-Sugar vs Control-Soda (blue), as well as (2b): Treatment-Soda vs Control-WH (orange) and Treatment-WH vs Control-Soda (blue). Bars represent mean percentages of mortality observed for each treatment group across the five mosquito species tested. Statistical differences in the percentages of mortality were assessed using Mann Whitney U tests, with asterisks (**) denoting statistical differences (*P*<0.01) in the orange vs. blue cage mean mortalities; error bars indicate SEMs.

Mortality rates were markedly higher in experimental cages containing Treatment-Soda baits. For instance, the Treatment-Soda mean mortality rate for *C. pipiens* (76.5 ± 14.65 %) was significantly higher than the corresponding mean mortality of 3.1 ± 1.93% observed in the Treatment-Sugar cages (P<0.01; Fig. 2a, S2 Table; corresponding negative ASB control data are shown in S3 Table). The highest mean percentage mortality for Treatment-Sugar trials across the four species was observed for *A. gambiae* (51.9% ± 15.74%), yet the corresponding mean mortality observed in the Treatment-Soda cages, with 100% mean mortality, was nearly twice as high (P<0.01; Fig 2a, S2 Table), correlating with a consistent preference for the soda baits.

Next the Soda and WH baits were compared, with the bait pairs including Treatment-Soda and Control-WH in one cage and Control-Soda and Treatment-WH in another. Once again, all mosquito species predominantly fed on the Treatment-Soda baits, as exhibited by the significantly higher mortalities in the Treatment-Soda groups compared to the Treatment-WH groups (Fig 2b, S4 Table; negative ASB control data are shown in S5 Table). For example, in *A. albopictus*, 84.8 ± 7.33% mosquitoes died in the Treatment-Soda cage, but only 16.0 ± 5.37% mortality was observed for the Treatment-WH cage (P<0.01; Fig 2b, S4 Table), the highest mortality observed in any Treatment-WH species trial. Conversely, the highest Treatment-Soda trial mortality was observed in *C. pipiens* (97.2 ± 1.38%) with only 7.0 ± 3.85% in corresponding Treatment-WH cages (P<0.01; Fig 2b, S4 Table). These results correlate with clear preferences for the soda baits.

### Evaluation of RNAi yeast-soda ATSB performance in semi-field trials conducted in the presence of competing natural sugar sources in Trinidad

To further assess the performance of the Sh.463_56.10R active ingredient in soda in a competitive tropical semi-field setting in Trinidad, the impact of the presence of competing natural sugar sources was evaluated by placing a flower or a piece of fruit in the Treatment-Soda mosquito cages. The mosquito strains used in these studies were generated from locally collected field specimens. The flowers assessed in these studies included *A. cathartica* (Fig. 3a,b), *C. roseus* (Fig. 3c,d), *H. rosa-sinensis* (Fig. 3e,f), *I. coccinea* (Fig. 3g,h), and *L. camara* (Fig. 3i,j), while the fruits included *C. papaya* (Fig. 3k,l) and *M. indica* (Fig. 3m,n), all of which are abundantly present in Trinidad. The results obtained when the mosquitoes were provided with a choice of the Treatment-Soda sachet and the fruit or flower were compared to those obtained in cages containing the Treatment-Soda sachets alone, with male and female preferences based on mortality recorded separately (Fig 3, S6 Table).

**Fig 3.**
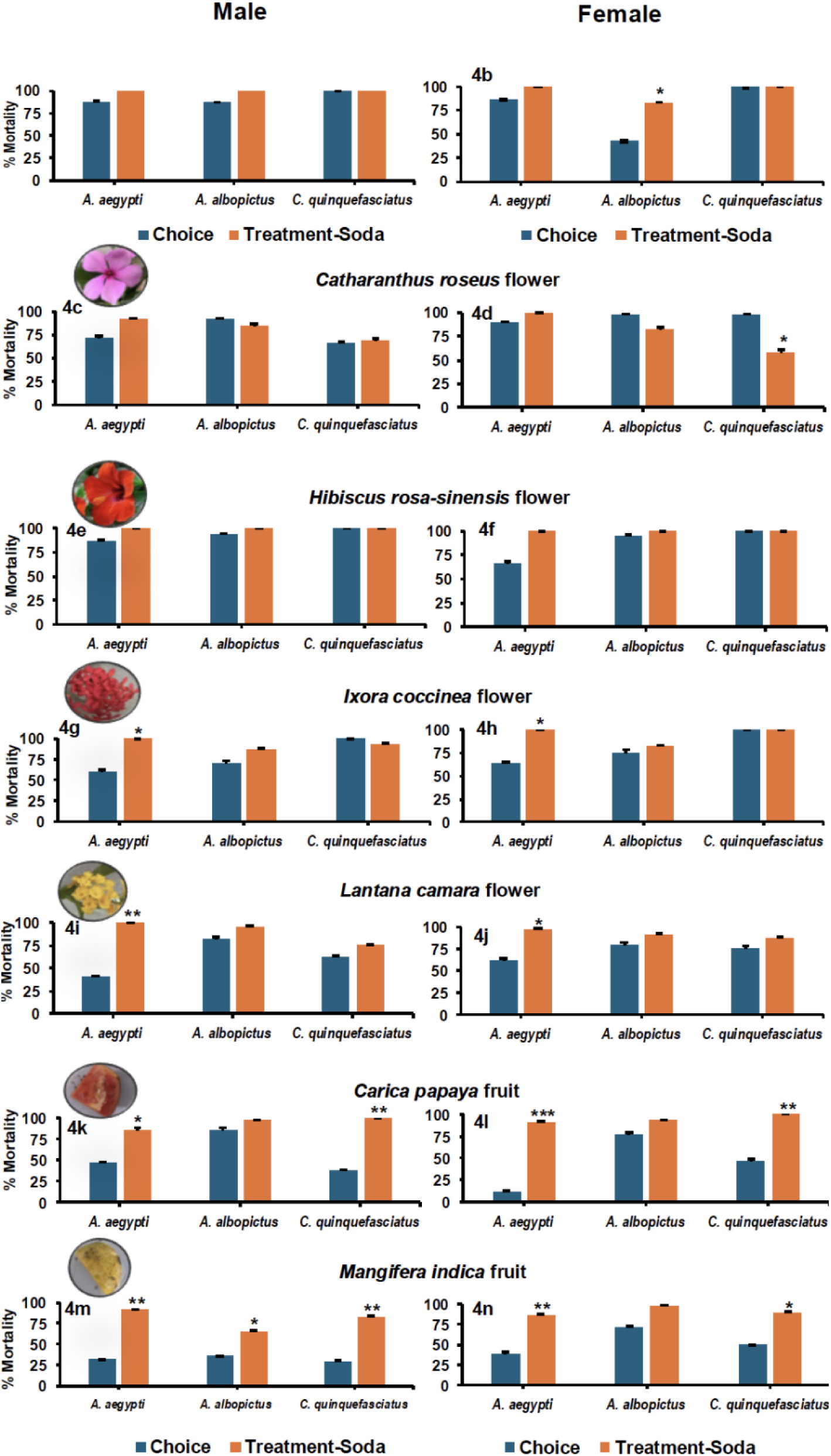
Semi-field assays conducted in Trinidad by gender. Results compiled from three replicate cage trials, each containing 15 male and 15 female mosquitoes were presented with Treatment-Soda sachets alone or in the presence of various fruits or flowers are shown. The results are reported separately for males and females of each of the indicated species. Statistical differences between the Treatment-Soda alone (orange) or in the presence of the indicated flowers/fruits (blue) are marked by asterisks (*= *P*<0.05, **= *P*<0.01, ***= *P*<0.001), with error bars indicating SEMs. (Fisher’s Exact Test).

For *A. aegypti* males and females, significantly less mortality (S6 Table) was observed in the presence of *I. coccinea* (P<0.05, Fig. 3g,h)*, L. camara* (P<0.01, Fig. 3i,j)*, C. papaya* (P<0.05, Fig. 3k,l), and *M. indica* (P<0.01; Fig. 3m,n), but mosquito mortality levels were not significantly impacted by *A. cathartica* (Fig. 3a,b)*, C. roseus* (Fig. 3b,c), or *H. rosa* (Fig. 3e,f) (S6 Table). Fewer impacts were observed in *A. albopictus,* with only *A. cathartica* significantly (P<0.05) reducing mortality rates in females (Fig. 3b) and *M. indica* significantly (P<0.05) reducing mortality rates in males (Fig. 3m) (S6 Table). *C. papaya* and *M. indica* fruits reduced *C. quinquefasciatus* mortality levels in both males (P<0.01) and females (P<0.05; Fig. 3k,l,m,n, S6 Table). It should be noted that in parallel to these trials, it was confirmed that the mosquitoes were able to feed on these fruits/flowers, as evidenced by observation of such feedings as well as generally low mortality rates recorded for mosquitoes provided with only the fruit/flower alone (or the result would have been death, which was not typically observed, S7 Table).

### Semi-field evaluation of a soda bottle feeder for RNAi yeast-soda ATSBs

A prototype soda bottle feeder was assessed in semi-field outdoor conditions in South Bend, IN. These trials were conducted on a *C. pipiens* strain established from a local collection. Mosquitoes were observed landing on and feeding from the soda feeders containing ASB-Soda, Control-Soda and Treatment-Soda feeders. Mosquitoes in the Treatment-Soda cages showed high mortality (94.7 ± 4.00%), dying within six days (Fig. 4). The mean percentage mortality of *C. pipiens* in the ASB-Soda (9.3 ± 6.73%) and Control-Soda (8.6 ± 6.70%) cages were significantly lower (*P* < 0.001) than the treatment cages (Fig. 4).

**Fig 4.**
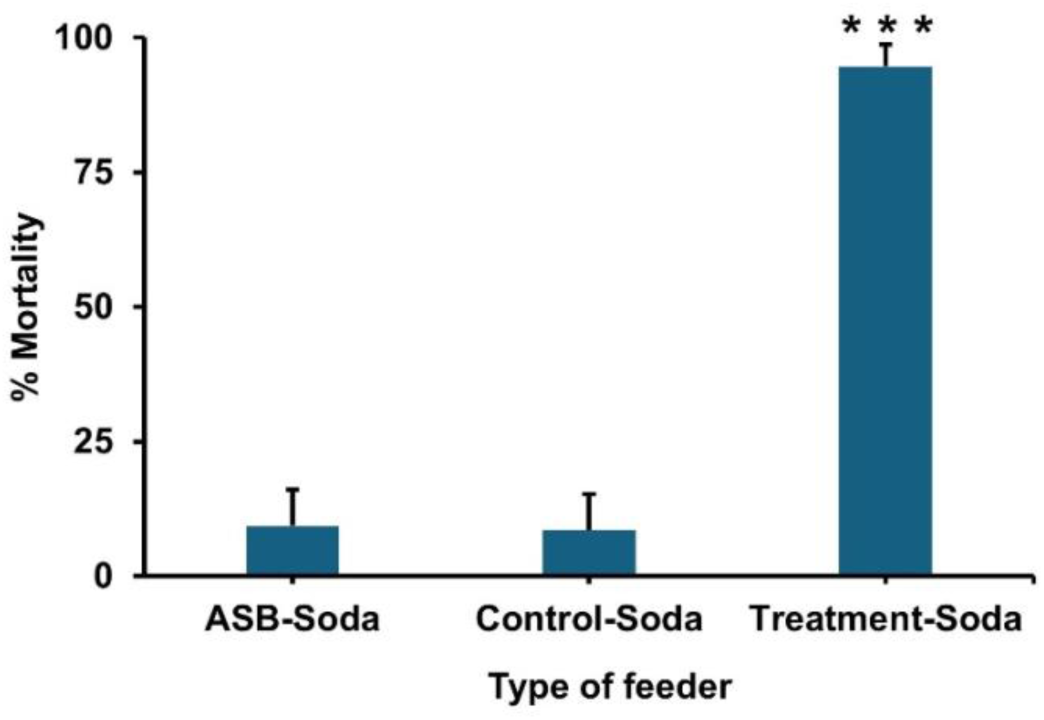
*C. pipiens* mortality in semi-field trials using soda bottle feeders. Significant mortality (*P*<0.001, denoted by ***) was observed in Treatment-Soda feeders when compared to ASB-Soda or Control-Soda (one-way ANOVA, Tukey post-hoc test). Results were compiled from nine replicate cages.

### Residual Activity of RNAi Yeast in Soda Over 14 Months

Residual activity of the Sh.463_56.10R Treatment-Soda bait was evaluated in laboratory trials conducted over the course of 14 months. The Treatment-Soda ATSB and Control-Soda bait were stored in the insectary to simulate field conditions for the duration of the experimental period. Throughout the trial, and across four different mosquito species evaluated, the Treatment-Soda retained high insecticidal activity in ATSB trials conducted intermittently on *A. aegypti, A. gambiae, A. stephensi and C. quinquefasciatus* (Fig. 5). Overall, monthly mean mortality rates ranged from 93-100% in the Treatment-Soda groups, while mean mortality rates of 0-14% were recorded in the Control-Soda cages across all species and timepoints. For each species, the Treatment-Soda vs. Control-Soda mean mortality levels throughout the fourteen-month trial period were significantly different (*P*<0.001, Fisher’s Exact test; Fig 5 and S8 Table).

**Fig 5.**
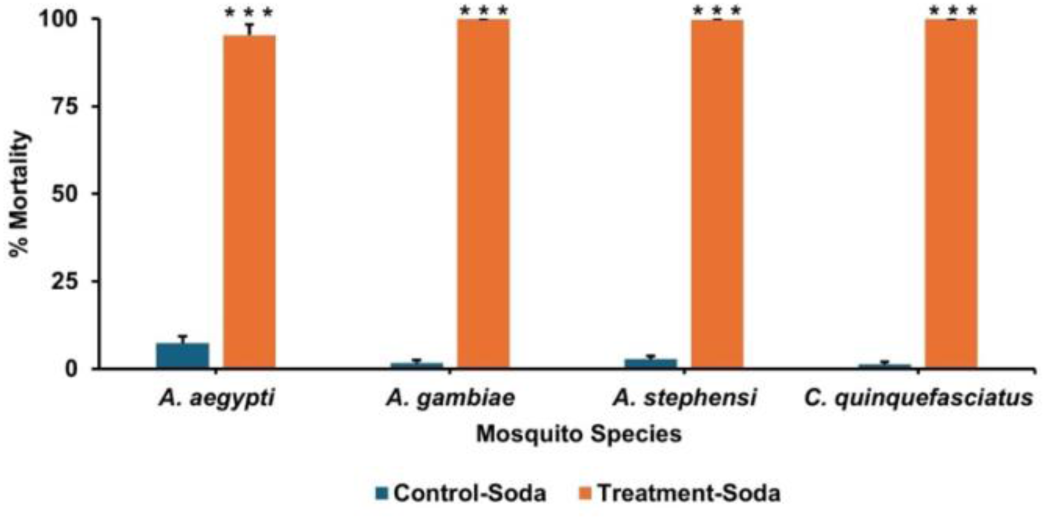
14 month residual activity of RNAi yeast in soda. The residual insecticidal activity of Sh.463_56.10R Treatment-Soda against four mosquito species tested across 14 months is shown. Mean mortality rates documented for *C. quinquefasciatus*, *A. aegypti*, *A. gambiae* and *A. stephensi* (n = 2 – 6 trials) following exposure to the Treatment-Soda and Control-Soda feeders. Bars represent mean mortality rates, with Control-Soda mortality consistently low across all species. Error bars indicate SEM. *** denotes significant differences (*P*<0.001) in mortality between the treatment groups (Fisher’s Exact test).

*C. quinquefasciatus* displayed 100% mortality during months 2-14 (S8 Table). Similarly, in *A. aegypti*, complete mortality (100%) was recorded at months 4 and 13, and a mortality of 93.0 ± 4.12% was recorded at month 10. Moreover, the Control-Soda groups consistently showed minimal mortality (0–10%). *A. gambiae* was evaluated at months 10 and 14, and the species exhibited a high susceptibility, with a mortality of 100% observed in the Treatment-Soda groups and Control-Soda mortality rates of 0.7 ± 0.67% and 4.5 ± 1.50%, respectively (S8 Table). *A. stephensi* also showed high susceptibility during the five timepoints it was tested with the bait. Mortality in the Treatment-Soda group ranged from 99.3 ± 1.50% in month 1 to 100% in the other four months, while mortality in the Control-Soda group remained low (0.0 ± 0.0% – 6.0 ± 2.00%) (S8 Table). For every ATSB trial conducted on each of the four species during the 14-month period, the Treatment-Soda vs. Control-Soda mean mortality levels were significantly different (*P*<0.001, S8 Table).

## Discussion

Maximizing the effectiveness of ATSBs as a scalable and environmentally sustainable vector control strategy requires optimizing their capacity to induce feeding in target mosquito populations (33). This depends on the effective integration of both the attractant and sugar components, which work together to lure mosquitoes to the baits, drawing them away from natural sugar sources, and ensuring ingestion of the toxic insecticide during sugar feeding (37). Enhancing ATSB delivery mechanisms not only improves kill rates but also extends coverage to a wider range of ecological settings and mosquito species (38).

Building on the importance of optimizing ATSB technology for mosquito control (38, 39), we showed that the active ingredient Sh.463_56.10R yeast delivered in three sugar types: Coca-Cola^TM^ (Soda), 10% sucrose (Sugar), and BaitStab^TM^ Westham bait matrix (WH), achieved high mortality across five mosquito species: *A. aegypti*, *A. albopictus*, *A. gambiae*, *C. pipiens*, and *C. quinquefasciatus* (Fig. 1, S1 Table). Mortality rates in the RNAi treatment groups consistently exceeded 75%, while mortality in the control yeast and ASB-only groups were negligible. These findings highlight the capacity of this yeast active ingredient to perform effectively across different sugar bait types, underscoring its versatility and operational flexibility for field applications (9, 11, 40). We have previously shown that Sh.463 and other RNAi-based mosquito-targeted pesticides can effectively induce species-specific mortality, thus reducing environmental risks (8, 9, 11, 40, 41). The oral delivery mechanism, specificity of the ATSB to target organisms, and scalable production of the yeast make this system well-suited for deployment in diverse ecological settings, particularly in areas where conventional interventions are failing due to resistance and behavioral avoidance (42).

Among the sugar formulations tested, soda emerged as the most effective vehicle for RNAi yeast delivery (Fig. 2, S2 Table). We observed a consistently strong feeding preference for RNAi yeast + soda baits, with mortality rates exceeding 93% across all species and reaching 100% for several Treatment-Soda trials. This suggests that beyond the intrinsic insecticidal activities of RNAi yeasts (8, 9, 11, 40, 41), components of soda (Coca-Cola^TM^) such as high sugar content, carbonation byproducts, or flavorings, may enhance phagostimulatory cues and palatability, thereby increasing mosquito feeding rates (35, 37, 43). This finding reinforces the superior attractiveness of soda-baited RNAi Treatments and their potential to outcompete natural sugar sources.

Previous studies have shown that incorporating fruit juices or fermenting sugars into ATSBs increases attractiveness and efficacy in *Anopheles* and *Aedes* mosquitoes (35, 44). Importantly, the increased mortality observed with lethal RNAi yeast + soda formulations in our study likely correlated with higher ingestion and delivery of RNAi-yeast load. These results support our hypothesis that combining RNAi yeast with a highly attractive sugar source can significantly enhance its entomological efficacy and operational viability as an ATSB-based mosquito control tool. Future studies should investigate the mechanisms underlying soda’s enhanced attraction, as well as explore its field-based validation.

Additionally, the potential synergistic effects between RNAi yeast insecticides and soda-based sugar formulations warrant further exploration to optimize their utility as ATSB tools. The efficacy of and high feeding preference for the RNAi yeast + soda ATSB in malaria vector species, *A. gambiae* and *A. stephensi,* demonstrated here and in a recent study (45) are also noteworthy, as malaria vectors are increasingly resistant to conventional control methods (3). Reports from recent large-scale ATSB field studies (29, 46), which tested ATSB technologies against *A. gambiae,* highlight the need for a more effective, scalable, user-friendly and environmentally sustainable ATSB tool. The RNAi yeast-soda bait evaluated in our study presents a promising platform to meet these criteria and advance ATSB-based mosquito control strategies.

The semi-field RNAi yeast + soda assays performed in the presence of sugars or fruits provide insight into how well this formulation might perform in the field (Fig. 3, S6 Table). These studies suggest that placing RNAi yeast-soda ASTB feeders near human dwellings requiring mosquito control could result in mitigating mosquito densities alongside the use of other mosquito control interventions, but that further investigation in the field is warranted. This is particularly the case for *A. aegypti,* which was impacted by a variety of different fruits and flowers (Fig. 3). The observed impacts of *L. camara* flowers (Fig. 3i,j, S6 Table) is consistent with observations made in the study by Upshur and colleagues (35) in which this flower was believed to be attractive to mosquitoes as a plant-sugar source, likely due to the presence of volatile benzaldehydes.

In terms of fruits, we observed all three mosquito species feeding on both *Carica papaya* and *Mangifera indica*, and the impacts of these fruits on RNAi yeast + soda efficacy in *A. aegypt, A. albopictus,* and *C. quinquefasciatus* are of potential concern (Fig. 3k,l,m,n, S6 Table). These fruits are both accessible and abundant during the fruit maturation period which often overlaps with the wet season in Trinidad from June to December. Field trials would help to better assess the competitiveness of the yeast in backyard environments in the presence of ripe fruits, including those assessed here and potentially other species. Many variables would have to be considered, for instance positioning of the bait around dwellings may still be impactful, particularly if the number of feeders is optimized. Another consideration is how effective application of these yeast baits may be during the bearing seasons where these and other fruits could be a more important sugar source for mosquitoes. The significant differences observed for both *C. papaya* and *M. indica* also suggest that extracts from these fruits can be explored as an additive to further increase attraction to Treatment-Soda bait stations.

The residual insecticidal activity of the Sh.463_56.10R Treatment-Soda bait was particularly striking. When stored under simulated field conditions in the insectary and tested monthly over a 14-month period, the Sh.463_56.10R Treatment-Soda bait continually induced high lethality in the four mosquito species tested (Fig. 5). *C. quinquefasciatus* exhibited consistently high mortality throughout the study period, with 100% mortality recorded in 11 of 12 trials and 98.5% in one month (S8 Table). Other species, namely *A. aegypti*, *A. gambiae*, and *A. stephensi* also responded strongly (monthly mortality rates: 93–100%) at all timepoints tested, including at month 14 (S8 Table). This is particularly notable given that control of arboviral vectors like *A. aegypti* and *C. quinquefasciatus* has historically relied on repeated insecticide applications and is often hindered by resistance (47, 48). The sustained RNAi-induced lethality observed here suggests a viable, long-lasting alternative to conventional adulticidal strategies for these species. Similarly, the residual effect against the malaria vectors, *A. gambiae* and *A. stephensi*, is important, and confirms the potential of RNAi yeast-soda ATSBs as a complementary tool for malaria vector control. This is especially relevant in the context of residual malaria transmission insecticide resistance and the recent spread of the invasive vector, *A. stephensi*, in urban areas of Africa (3, 49). Overall, these results demonstrate that inactivated RNAi yeast remains stable and bioactive over extended periods, even under simulated field conditions, a critical property for products intended for deployment in tropical environments. Importantly, our findings support the feasibility of soda-baited RNAi treatments as an effective, long-lasting and universally attractive mosquito control tools.

Finally, it should be noted that although the soda bottle feeders performed well (Fig. 4), they were difficult to construct, and leaking could sometimes be observed. This may be due to the lower surface tension of soda (approximately 42.2 mN/m) compared to sugar water (73.5 mN/m) (50). The design worked well enough for a prototype, but simplifying construction and correcting the leakage would be a helpful and seemingly straightforward next step, as an easy to construct low-maintenance feeder would be ideal.

## Conclusions

This study has demonstrated the potential for using soda-baited RNAi yeast as a potent and scalable platform for deploying ATSB-based mosquito control. Its ability to induce high mortality across multiple species, combined with long-term residual activity, makes it a promising candidate for field-based implementation, particularly in resource-limited settings and as part of integrated vector management programs. Additionally, yeast-based formulations offer a cost-effective, environmentally friendly, species-specific alternative to chemical insecticides. The semi-field trials in Trinidad demonstrated that the RNAi yeast-soda bait may be competitive against some natural sugar sources. Future research should focus on evaluating the bait’s operational performance under field conditions in small-scale, and eventually in large-scale field trials.

## Acknowledgements

Thanks to members of our labs for their useful suggestions, particularly Keshav Mysore for his meaningful input. We also thank our UWI colleagues for their help and advice.

## Supporting Information Captions

**S1 Table: Mortality rates of various mosquito species exposed to RNAi yeast in three different sugar baits.** Mosquito mean mortality rates following exposure to the different RNAi yeast-sugar formulations in laboratory trials. Mortality across the five mosquito species is expressed as mean ± SEM (%). The number of female mosquitoes tested was N=225. P-values were calculated using the one-way Kruskal-Wallis-Test to compare mortality among Treatment (Sh.463_56.10R), Control (Control_347.1R) and ASB (Sugar base only) groups for each formulation. Dunn’s post hoc tests were used to determine which comparisons were significantly different (P<0.01).

**S2 Table: Mosquito mortality rates for Treatment-Soda and Treatment-Sugar RNAi yeast baits.** Mosquito mean mortality rates following exposure to the Treatment-Soda (vs Control-Sugar) and Treatment-Sugar (vs Control-Soda) RNAi yeast-sugar formulations in laboratory trials. Mortality across the five mosquito species, with a total of N=150 females per group, is expressed as mean ± SEM (%). Mann Whitney U tests were used to determine statistical differences (P<0.01) in mortality between Treatment-Soda and Treatment-Sugar experiments.

**S3 Table: Mosquito mortality rates for ASB-Soda and ASB-Sugar baits.** Mosquito mean mortality rates for control experiments, which were conducted in independent cages alongside dual choice experiments described in S2 Table. Mortality across the five mosquito species, N=150 females per group, is expressed as mean ± SEM (%).

**S4 Table: Mosquito mortality rates for Treatment-Soda and Treatment-WH RNAi yeast baits.** Mosquito mean mortality rates following exposure to the Treatment-Soda (vs. Control-WH) and Treatment-WH (vs. Control-Soda) RNAi yeast-sugar formulations in laboratory trials. Mortality across the five mosquito species, n=3 with a total of N=150 females per group, is expressed as mean ± SEM (%). Mann Whitney U tests were used to determine statistical differences (*P*<0.01) for the mortality between Treatment-Soda and Treatment-WH experiments.

**S5 Table: Mosquito mortality rates for ASB-Soda and ASB-WH baits.** Mosquito mean mortality rates for control experiments which were conducted in independent cages alongside the experiments described in S4 Table. Mortality across the five mosquito species, N=75 females per group, is expressed as mean ± SEM (%).

**S6 Table: Semi-field choice trials conducted using RNAi yeast baits in the presence of competing natural sugar sources in Trinidad.** Results from ATSB feeding assays conducted on male (n=15) and female (n=15) mosquitoes of the specified species. Mosquitoes were presented with a choice of the indicated fruits/flowers and a Treatment-Soda sachet. Statistical differences (Fisher’s Exact Test, P<0.001) for triplicate cage trials (N=45) is expressed as mean ± SEM (%).

**S7 Table: Semi-field trials with natural sugar sources conducted on three mosquito species in Trinidad.** Results from feeding assays conducted on male (n=15) and female (n=15) mosquitoes for three replicate cage trials (N=45) with the indicated species that were provided with only the specified flowers or fruits. Average percentage mortality for triplicate experiments is expressed as mean ± SEM (%).

**S8 Table: Residual activity of RNAi yeast in soda over 14 months.** Residual activity of the RNAi yeast in soda was evaluated in the laboratory over a 14-month period. Mortality rates (mean %) for n = 2 – 6 trials are presented for Treatment-Soda and Control-Soda across the different mosquito species and experimental time points. Significant differences (P<0.001) in mortality between Control-Sugar and Treatment-Soda for the specified species in the indicated months are shown (Fisher’s Exact tests).

## Notes

### Competing Interest Statement

MDS and DWS are inventors of U.S. patent number 62/361,704 and European application number 17828458.4.

